## Supplementary text for "Theoretical model of membrane protrusions driven by curved active proteins"

### Supplementary Material

#### S-1. CALCULATION OF THE PERIMETER OF CMC CLUSTERS

The CMC-bare membrane boundary is measured by summing the dual of the edges between the cluster and bare membrane. These are the edges in the voronoi lattice, connecting the mid-section of each edge to the circumcenter of the adjacent triangles i.e. the center of the inscribing circle (see figure S-1). Partitioning each triangle between its vertices is already used in the calculation of the curvature [1].

#### S-2. ANALYTICAL CALCULATION OF THE FLAT-PEARLING PHASE TRANSITION LINE

We can make a rough analytical estimation for the flat-pearling transition by equating the active work and energy of the flat phase from a mixed phase to the energy of the pearling phase (Fig. S-2). In the flat phase, moving the active CMCs outwards from the radius of the sphere $r_p$ to the larger radius of the flattened disc $r_f$ results in work. The pearling phase has binding advantage because all CMC vertices are connected, with $-w$ per edge, while the flat rim has large interface (boundary perimeter length) where CMCs vertices neighbor bare membrane vertices, whose edge does not contribute. The pearling phase has a bending disadvantage due to the bare membrane body, which is roughly spherical with an energy of $8\pi\kappa$, compare to the flat phase where the bare membrane is in two flat discs with no bending energy (both the pearling and rim clusters are curved to fit the CMCs, so they do not have bending energy).

$$-\left(r_f - r_p\right) F = -w\left(\chi_p - \chi_f\right) + 8\pi\kappa \tag{S-1}$$

The radius difference $\Delta r = r_f - r_p$ (Fig. S-2), and the number of CMC-CMC bonds $\chi_p, \chi_f$ in the pearling and flat phases respectively, are dependant on the geometry of the phases, so they should be very weakly dependant on the specific model parameters. Therefore $\Delta r$ and $\chi_p - \chi_f$ do not depend on $w, f, \kappa$, and we end up having a linear relation between $f$ and $w$ along the transition line in the $f, w$ phase diagram. In the force-binding strength $(f - w)$ system, we take the values for these geometric quantities from simulations and draw the resulting line on the phase diagram (Fig. S-3, green line), which qualitatively matches the behavior of the transition observed in the simulations.

#### S-3. MIXED CURVATURE CMC CLUSTERS

The concave and convex CMCs generate a wavelike pattern, but analyzing it in terms of wavenumber is difficult, since the clusters are part of an irregular, triangulated surface. The undulations of the CMCs in the mixed clusters are essentially independent of $C_0$, and $f$, as shown in Fig.S-4. Note that we are at the limit of the mesh resolution for these undulations. We have yet to be able to compare this to the experimental results in [2].

#### S-4. MIXED CURVATURE WITH EXCLUSIVE BINDING

The mixed curvature system (Fig. 4c in the main text) was also simulated using exclusive binding, i.e. only same-curvature CMCs bind together (Fig.S-5). The result is that the two CMCs types form separated aggregates, with the active convex CMCs aggregating along the rim and forming the flat phase. The passive concave CMC form separated clusters of different shapes, depending on their spontaneous curvature. Highly concave CMCs ($C_0^- \leq -0.45$) aggregate into internal pearling clusters, that do not affect the flat global phase. The shallower concave CMCs ($C_0^- \geq -0.3$) aggregate into large, shallow bowl-like patches.

In some cases, these concave aggregates are able to form with convex CMC along their rim, since their curvatures complement each other (see for example at $C_0^- = -0.3$). Since the convex active CMC along the rim of the concave cluster apply protrusive forces, they end up forming together a "cup"-like protrusion. When the force is inhibited, this aggregation occurs, but it is not elongated as a protrusion (compare "None" with "Disable" at $C_0^- = -0.3$ in Fig.S-5). Other than that, inhibition doesn't appear to significantly affect the results in Fig.S-5, since there is no significant contact between the two CMC types. These shapes, in the form of open bowls, resemble early stages of macropinocytosis [3, 4], but do not evolve to induce closure of the "mouth", as we observed when the convex and concave CMC had direct interactions (Fig.5 in the main text).

| parameter | units | Fig.S-3 | Fig.S-4 | Fig.S-5 | Fig.S-6 | movie 1,2 | movie 3,4 |
|---|---|---|---|---|---|---|---|
| $f$ | $K_BT/\ell_{min}$ | $0-1.2$ | $0.5, 0$ | 0.5 | 0.5 | 0.2, 0.5 | 0.5 |
| $w$ | $K_BT$ | $0-0.48$ | 2 | 2 | 2 | 2 | 2 |
| $\kappa$ | $K_BT$ | 20 | 28.5 | 28.5 | 28.5 | 28.5 | 28.5 |
| $\rho$ | 1 | 10% | $10\%, 10\%$ | $10\%, 10\%$ | $10\%, 10\%$ | 20% | $10\%, 10\%$ |
| $C_0$ | $1/\ell_{min}$ | 1 | -0.6 $-$ 0, 0.8 | -0.75 $-$ 0, 0.8 | 0.8 | 0.4,0.1 | 0.8 |

TABLE I: The values of the model parameters used in the simulations in the different figures. The energy units are $K_BT = 1$, which define the scale of $f, w, \kappa$, and the length units are $\ell_{\min} = 1$, which define the scale of the vertex lattice, the force, and spontaneous curvature.

#### S-5. VESICLES WITH BOTH NORMAL AND ALIGNED-FORCE CMC, ADHERED TO A FLAT SUBSTRATE

In Fig.S-6 we show the dynamics of the vesicle that contains the mixture of aligned-force (yellow) and normal-force (red) CMC, which have exclusive binding interactions between them (see Fig.8 in the main text). At time $t = 250$ we turned off the normal-force CMC, keeping only the aligned-force CMC active. We find that the adhered area shape changes, with the rim regions that contain the curved passive (red) CMC retract into the vesicle, while the aligned-force regions protrude more prominently along the adhered rim.

##### Movies

- **Movie-S1** Aligned-force simulation of the formation of filopodia-like tubular protrusions (corresponding to Fig.7B), with parameters $\kappa = 28.5, f = 0.2, w = 2, C_0 = 0.4, \rho = 20\%, s = 0.5, r = 30$
- **Movie-S2** Normal force simulation, in the regime of tubes shapes (corresponding to Fig.7B), with parameters $\kappa = 28.5, f = 0.5, w = 2, C_0 = 0.1, \rho = 20\%$
- **Movie-S3** Adhered, universal-binding between normal-force CMCs (red) and aligned-force CMCs (yellow), corresponding to Fig.8A. Parameters used: $\kappa = 28.5, f = 0.5, w = 2, w_{ad} = 0.25, C_0 = 0.8, \rho_n = 10\%, \rho_a = 10\%, s = 1, r = 15$
- **Movie-S4** Adhered, exclusive-binding between normal-force CMCs (red) and aligned-force CMCs (yellow), corresponding to Fig.8A. Parameters used: $\kappa = 28.5, f = 0.5, w = 2, w_{ad} = 0.25, C_0 = 0.8, \rho_n = 10\%, \rho_a = 10\%, s = 1, r = 15$
- **Movie-S5**. The 3D movie of the cell in Figure 9A
- **Movie-S6**. The movie of the XY and XZ section for Figure 9E
- **Movie-S7**. The movie of the XY and XZ section for Figure 9F
- **Movie-S8**. The movie of the XY and XZ section for Figure 9G


[1] G. Gompper and D. M. Kroll, in *Statistical Mechanics of Membranes and Surfaces* (WORLD SCIENTIFIC, 2004), pp. 359–426.

[2] E. Sitarska, S. D. Almeida, M. S. Beckwith, J. Stopp, Y. Schwab, M. Sixt, A. Kreshuk, A. Erzberger, and A. Diz-Muñoz, bioRxiv p. 2021.03.26.437199 (2021).

[3] D. M. Veltman, T. D. Williams, G. Bloomfield, B.-C. Chen, E. Betzig, R. H. Insall, and R. R. Kay, Elife **5**, e20085 (2016).

[4] R. R. Kay, Cells & Development **168**, 203713 (2021).

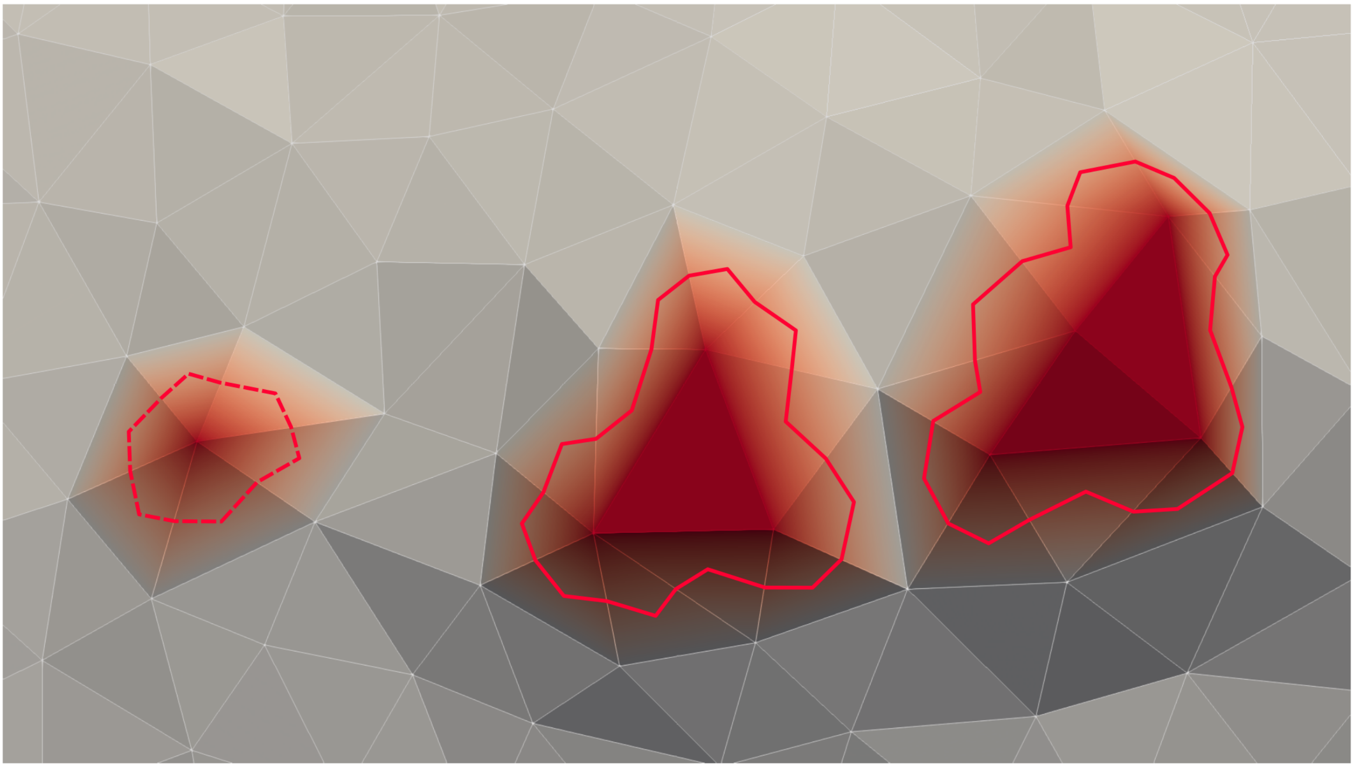

FIG. S-1: Sketch of the boundary of connected clusters: for each edge between the cluster and the outside, a line is drawn from the middle to the center each of the adjacent triangles. We ignore the single-clusters (dashed line)

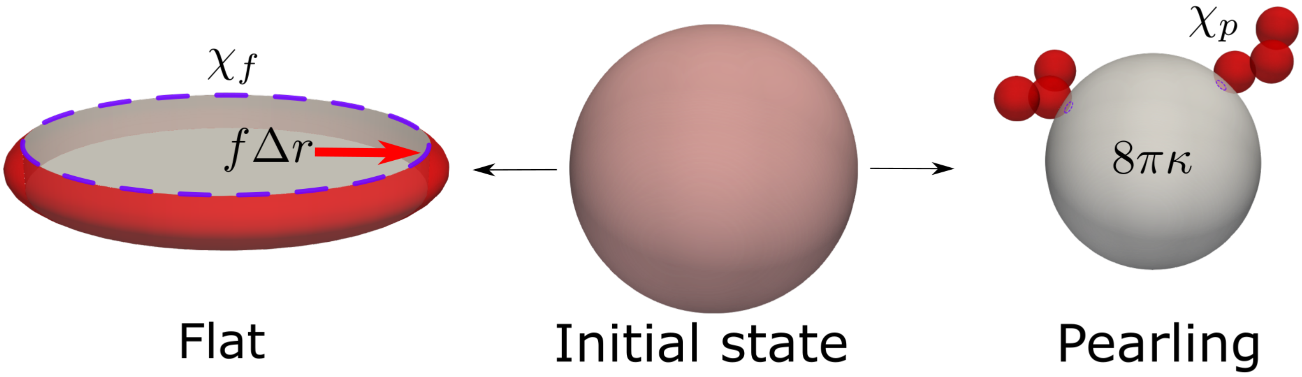


FIG. S-2: Schematic description of the transition between flat and pearling phases, from an initially mixed, spherical phase (at the center). Bare membrane is in white, and CMCs in red, and mixed composition in pink. The flat transition result in all CMCs moving from the surface of the sphere to the rim of a flat disc, which has a larger radius $\Delta r$. Due to active force $f$, this generates work $W = -f\Delta r$. The bending energy of the CMCs on the rim and in the pearling clusters is assumed to be approximately 0, but the spherical body of bare membrane in the pearling phase has a bending energy of a closed sphere: $8\pi\kappa$, while it is zero for the flat discs of bare membrane in the flat phase (since they are flat). Finally, the number of CMC-CMC bonds in the pearling phase $\chi_p$ is larger than in the flat phase $\chi_f$, since in the flat phase it is reduced due to the large boundary between the rim cluster the the flat bare membrane discs.

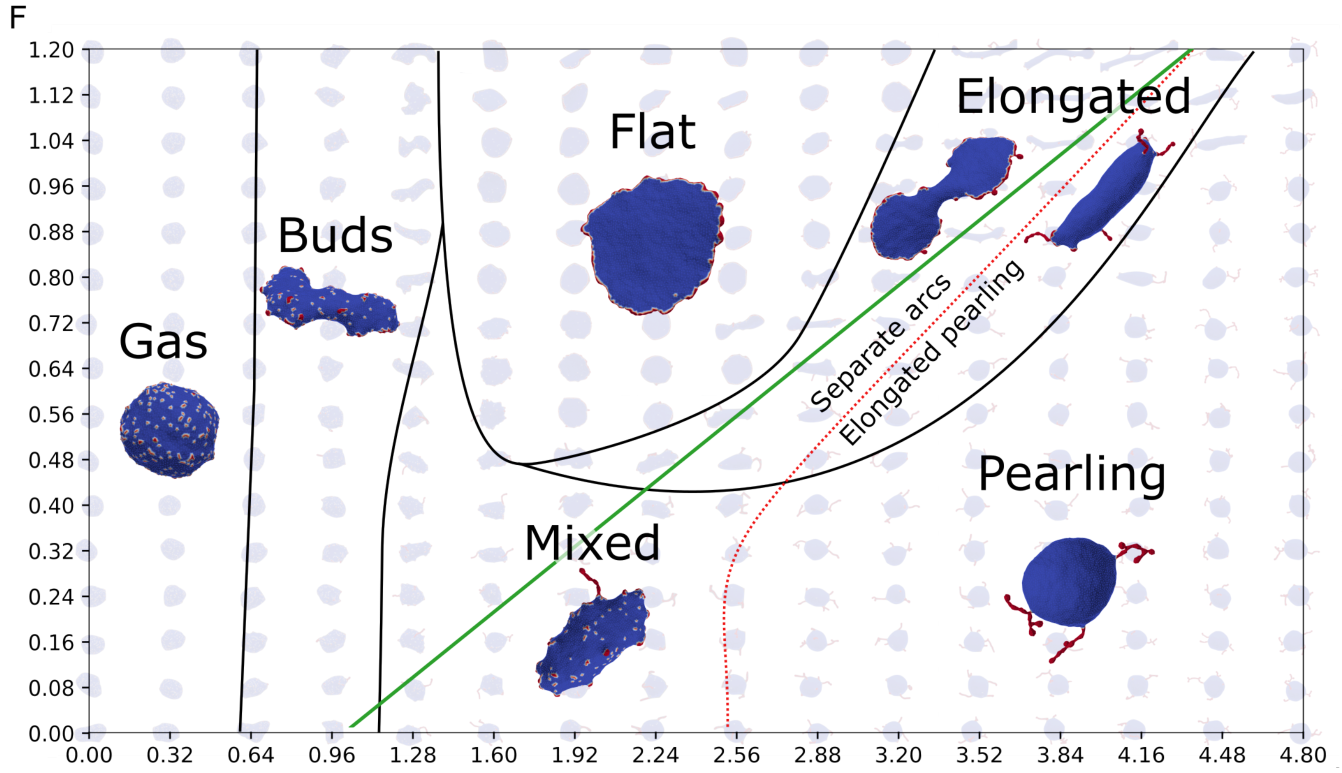


FIG. S-3: Phase diagram of the force-binding strength system, with an analytically-derived transition line for the pearling-flat transition (green line, Eq.S-1).

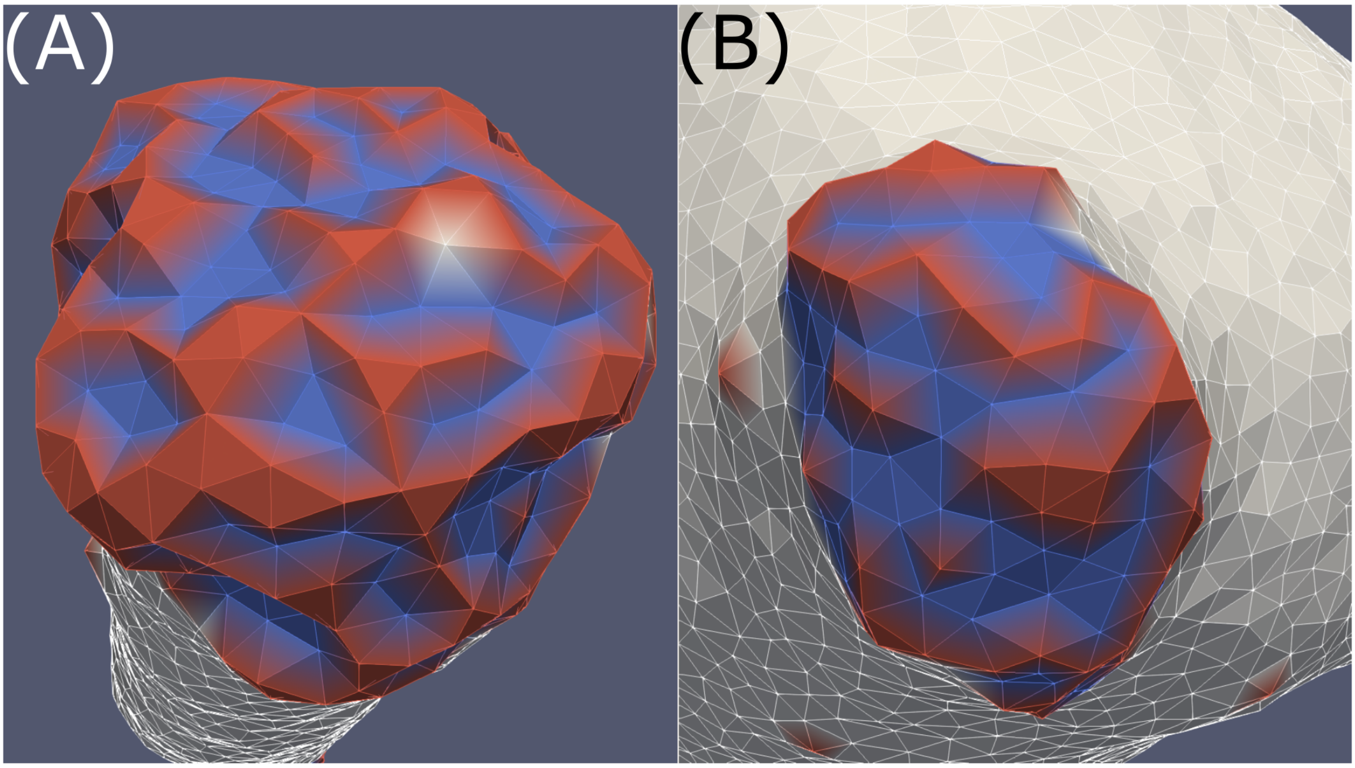


FIG. S-4: The undulation of a CMC cluster with (A) highly concave (−0.6) active CMC (B) with shallow concave (−0.001) CMC and disabled force. The size and shape of the clusters is very different, but the peaks and troughs patterning due to CMC shape is at the limit of the mesh resolution for both.

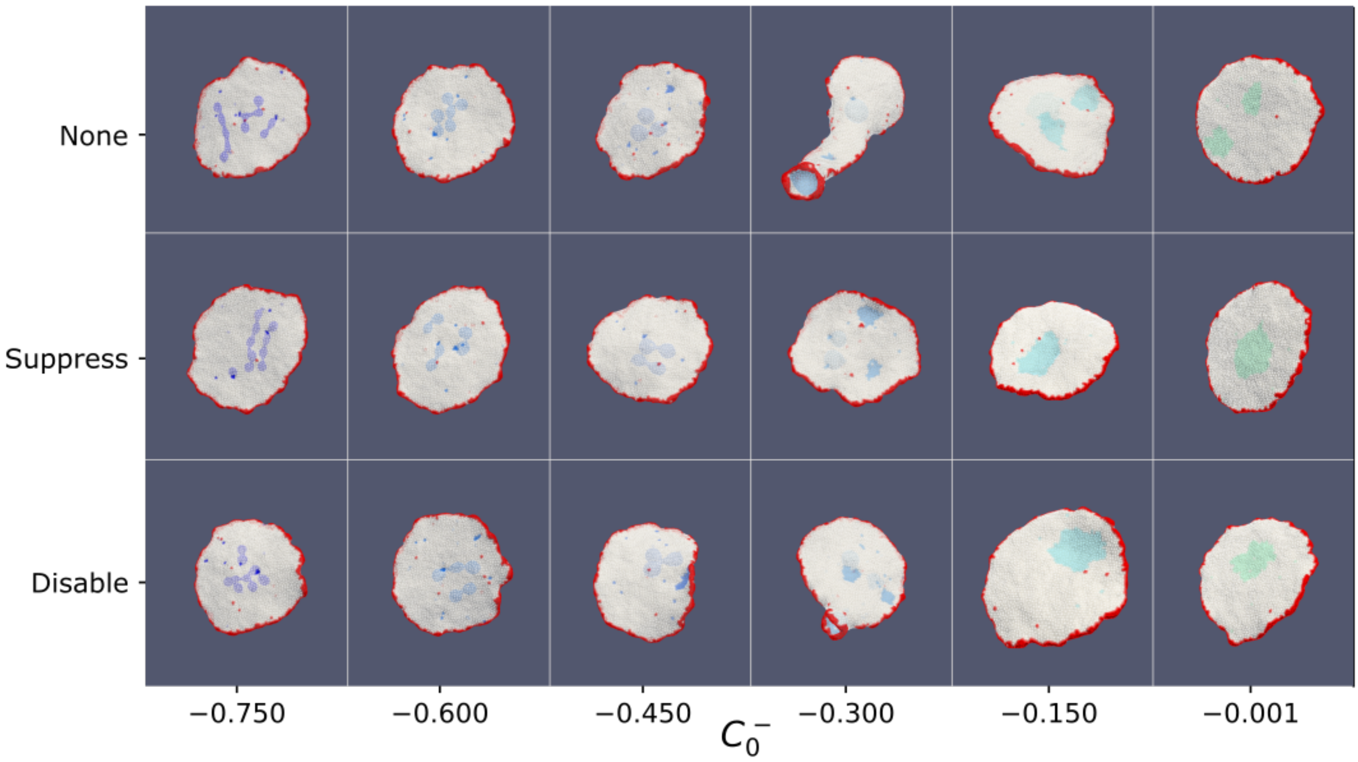


FIG. S-5: Active convex and passive concave system (red and blue, respectively), with binding between same type only. As in the universal binding case, the suppressive and disabling inhibition do not have any strong effects, since the types are separated. Simulations with $C_0^- \leq -0.3$ are draw semi-transparent. In all cases, the convex CMCs aggregate in a rim, making the vesicle flat, and concave CMCs aggregate in pearling for $C_0^- < -0.3$, bowl-like patches for $C_0^- > -0.3$, and both for $C_0^- = -0.3$.

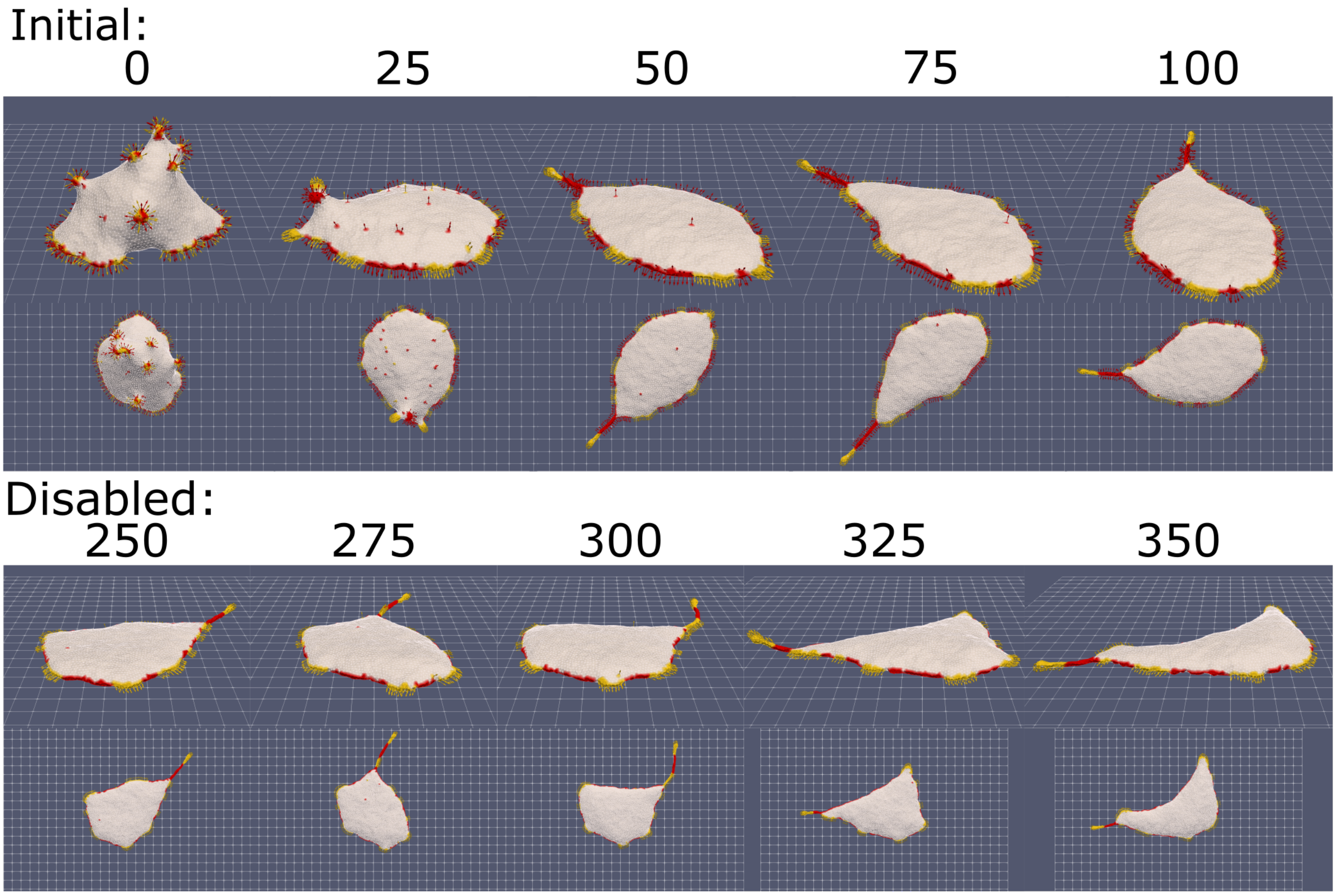


FIG. S-6: Overview of an adhered vesicle with a mixture of aligned-force (yellow) and normal-force (red) CMC, which have exclusive binding interactions between them (see Fig.8 in the main text). At time $t = 250$ the force is disabled for the normal-force CMCs, leaving only the aligned-force CMCs active. The original simulation is given on the top (times $0 - 100$), and the simulation after the normal-force has been disabled is at the bottom.
